# Copper acquisition is essential for plant colonization and virulence in a root-infecting vascular wilt fungus

**DOI:** 10.1101/2024.04.23.590778

**Authors:** Rafael Palos-Fernández, María Victoria Aguilar-Pontes, Gema Puebla-Planas, Harald Berger, Lena Studt-Reinhold, Joseph Strauss, Antonio Di Pietro, Manuel Sánchez López-Berges

## Abstract

Phytopathogenic fungi provoke devastating agricultural losses and are difficult to control. How fungal pathogens adapt to the plant environment to cause disease and complete their life cycle on the host remains poorly understood. Here we show that efficient acquisition of copper, mediated by the transcriptional regulator Mac1, is crucial for plant colonization and virulence in *Fusarium oxysporum,* a soilborne ascomycete that causes vascular wilt on more than 150 different crops. RNA-seq and ChIP-seq establish a direct role of Mac1 in activation of copper deficiency response genes, many of which are induced during plant infection. Loss of Mac1 impairs growth of *F. oxysporum* under copper-limiting condition as well as vascular colonization and virulence on tomato plants. Importantly, Mac1-independent overexpression of a copper reductase and a copper transporter restores growth under copper limitation and virulence in the *mac1* null mutant background. These findings establish a key role for copper acquisition in fungal pathogenicity on plants and reveal new ways to protect crops from phytopathogens.

## INTRODUCTION

Vascular wilt fungi constitute a particularly destructive group attacking almost every crop except cereals and are extremely difficult to control(Berendsen et al., 2012). The *Fusarium oxysporum* (*Fo*) species complex provokes devastating losses in global agriculture(Dean et al., 2012). Its destructive potential is exemplified by an aggressive clone named tropical race 4 (TR4) that is currently threatening the world’s most important staple crop banana(Ordonez et al., 2015). *Fo* infection initiates in the soil, when the fungus senses chemical signals released by roots that trigger directed hyphal growth towards the plant(Turra et al., 2015). After entering the plant, *Fo* initially grows intercellularly in the root cortex and subsequently enters and colonizes the xylem vessels, causing characteristic wilt symptoms and plant death(Dean et al., 2012). Besides provoking wilt disease in plants, *Fo* is also an opportunistic pathogen of humans causing symptoms ranging from superficial skin and cornea infections to lethal systemic fusariosis(Nucci and Anaissie, 2007).

During the infection process, fungal pathogens compete with the host for limited nutrients and microelements. The latter include transition elements such as iron, copper or zinc, which act as essential cofactors for key cellular processes like electron transfer. All fungi have evolved mechanisms to ensure the efficient uptake and use of these metals under limiting conditions(Rutherford and Bird, 2004). For example, adaptation to iron limitation was previously shown to be critical for fungal virulence on both plant and animal hosts(Schrettl et al., 2010; López-Berges et al., 2012).

Similar to iron, copper exists in two relevant oxidation states, Cu^+^ and Cu^2+^, acting as a cofactor for enzymes due to its potential to either accept or donate an electron while switching between the two states(Smith et al., 2017). Moreover, copper can bind to certain proteins thereby stabilizing their conformation(Festa and Thiele, 2011). In the model fungus *Saccharomyces cerevisiae*, adaptation to copper limitation is mediated by the transcription factor Mac1(Wegner et al., 2011; Shi et al., 2021), which is conserved in most filamentous fungi(Wiemann et al., 2017; Raffa et al., 2019). Under conditions of copper deficiency (-Cu), Mac1 directly binds to and transcriptionally activates copper uptake genes such as those encoding metalloreductases that convert Cu^2+^ into Cu^+^, or high-affinity copper transporters that internalize Cu^+^(Smith et al., 2017; Raffa et al., 2019). Mac1-mediated adaptation to copper limiting conditions was previously shown to be important for virulence in human fungal pathogens such as *Aspergillus fumigatus*, *Cryptococcus neoformans*, *Histoplasma capsulatum* or *Candida albicans*, which face copper limitation in certain host tissues such as kidney or brain(Waterman et al., 2007; Sun et al., 2014; Cai et al., 2017; Raffa et al., 2019; Culbertson et al., 2020; Ray and Rappleye, 2022).

The role of copper uptake during fungal infection of plant hosts has not been explored so far. Here we show that -Cu response genes such as metalloreductases and high-affinity copper transporters are markedly upregulated in *F. oxysporum* during colonization of tomato roots and that this upregulation is directly mediated by Mac1. We further demonstrate that reduction of Cu^2+^ to Cu^+^ and its subsequent uptake by the fungal cell are essential for *Fo* virulence. Our results reveal a previously unknown role of copper uptake in fungal pathogenicity on plants and suggest novel ways to control plant disease.

## RESULTS

### *F. oxysporum* Mac1 is essential for adaptation to copper limiting conditions

A BLASTp search of the genome database of *Fusarium oxysporum* f. sp. *lycopersici* 4287 (*Fol4287*) using the Mac1 amino acid sequences of *S. cerevisiae* (YMR021C)(Jungmann et al., 1993) and *A. fumigatus* (Afu1g13190)(Park et al., 2017) identified a single putative Mac1 ortholog, FOXG_03227, with a predicted CDS starting at position 1,808,878 of chromosome 8 (NC_030993.1). Initial inspection revealed that the protein predicted in the database was significantly shorter than the homologs from *S. cerevisiae* and *A. fumigatus* and lacked the copper-fist DNA-binding domain. Manual inspection of the DNA sequence surrounding *FOXG_03227* identified an additional putative ATG start codon at position 1,808,411, giving rise to a CDS of 1,607 bp and an intron between position 33 and 103. The predicted protein has a length of 511 aa, shows 26.92% and 30.02% identity with *S. cerevisiae* and *A. fumigatus* Mac1 proteins, respectively, and contains the conserved Mac1 copper-fist DNA-binding and Cu-binding domains (Figure S1A and B). We therefore concluded that the newly annotated *FOXG_03227* ORF corresponds to the correct *mac1* gene of *Fol4287*. Phylogenetic analysis with characterized Mac1 proteins from different fungal species revealed that *A. fumigatus* Mac1 is the closest ortholog and that the orthologs of *C. neoformans* and *S. pombe* are closer to *Fo* Mac1 than those of *C. albicans* and *S. cerevisiae* (Figure S1C). A *mac1*Δ mutant was generated by replacing the complete *mac1* ORF in *Fo* with the *Hyg*^R^ resistance cassette (Figure S2A and B). The *mac1*Δ strain was subsequently complemented in locus with a DNA construct containing the wild-type *mac1* ORF fused at the 3’ end to the S-tag oligopeptide (*mac1*^Stag^) (Figure S2C and D).

To determine the role of Mac1 in adaptation of *Fo* to -Cu conditions, growth rate on solid medium and biomass production in liquid media of the *mac1*Δ mutant were compared to those of the wild-type and the *mac1*^Stag^ complemented strain. Growth of the *mac1*Δ mutant was drastically impaired on -Cu solid media but was similar to that of the wild-type strain at CuSO_4_ concentrations starting at 10 µM, including the toxic concentration of 2 mM CuSO_4_ (Figure 1A). The complemented *mac1*^Stag^ strain displayed the same phenotype as the wild-type suggesting that the C-terminal S-tag fusion of Mac1 is fully functional. Biomass production of the *mac1*Δ mutant in -Cu was significantly reduced compared to the wild-type and the complemented strain. Unexpectedly, *mac1*Δ produced significantly more fungal biomass than the wild-type or *mac1*^Stag^ strains when grown in the presence of 10 µM CuSO_4_ (Figure 1B).

**Figure 1.**
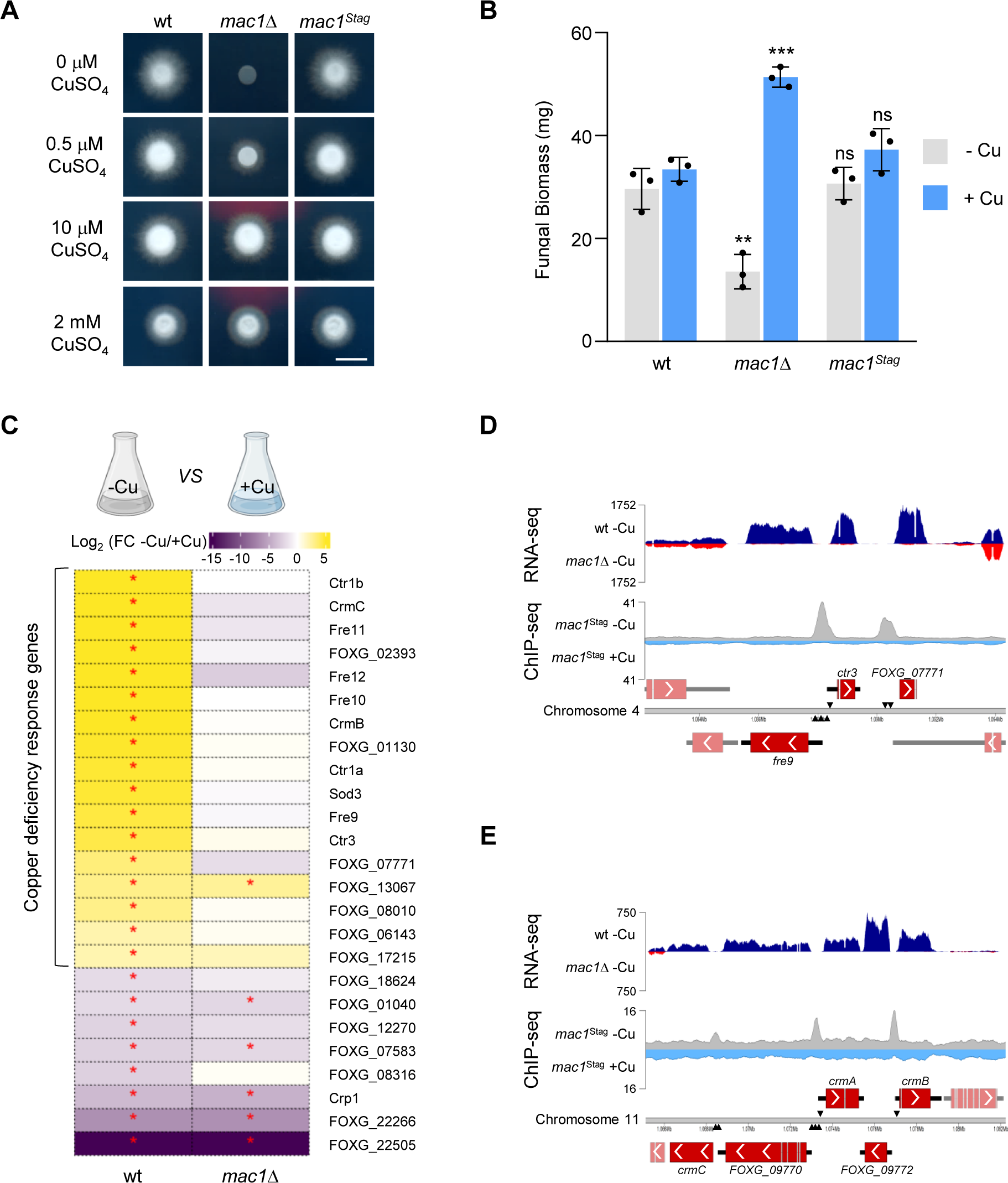
*Fusarium oxysporum* Mac1 transcriptionally activates copper deficiency response genes and is required for adaptation to copper limitation. **A.** Mac1 is required for growth under copper-limiting conditions. Colony phenotypes of the wild type (wt), the deletion mutant (*mac1*Δ) and the *in-locus* complemented strain (*mac1*^Stag^) after 2 d growth on minimal medium containing 20 mM L-glutamine, pH 6.5 and trace elements lacking copper (MM+TE^-Cu^), supplemented with the indicated concentrations of CuSO_4_. Scale bar, 1 cm. **B.** Fungal biomass (dry weight) obtained from the indicated strains after 16 h gemination in MM+TE^-Cu^ supplemented (+Cu) or not (-Cu) with 10 µM CuSO_4_. Bars represent standard deviations (*n* = 3, biological replicates). *p*-values: ns>0.05, **<0.01, ***<0.001 versus wt under the same condition according to two-tailed unpaired Student’s *t* test. **C.** Fold change (FC) of transcript levels of the indicated genes in the wt (left column) and the *mac1*Δ strain (right column) under -Cu versus +Cu conditions was measured by RNA-seq. Strains germinated 15 h at 28 °C in Potato Dextrose Broth were transferred for 6 additional h to MM+TE^-Cu^ with (+Cu) or without (-Cu) 100 µM CuSO_4_. Differentially expressed genes are ordered according to FC in the wt. *p*-value: *≤0.05 within each comparison. Data were calculated from three independent biological replicates. **D, E.** Abundance of RNA-seq transcript reads of the wt (dark blue) or the *mac1*Δ strain (red) under -Cu conditions (RNA-seq, upper graphs); or of gDNA reads from ChIP-seq analysis in the *mac1*^Stag^ strain under -Cu (grey) or +Cu (light blue) conditions (ChIP-seq, lower graphs). Data are represented as base-level coverage to two *Fol4287* gene clusters, harboring the *ctr3*, *fre9* and *FOXG_07771* genes (D) or the *crmC*, *FOXG_09770*, *crmA*, *FOXG_09772*, and *crmB* genes (E). Genes are indicated as red boxes and putative Mac1 binding sites on each strand by black triangles.

### Mac1 directly activates expression of copper limitation response genes

We next determined the role of Mac1 in the transcriptional response of *Fo* to -Cu conditions. RNA-seq analysis of the wild-type strain grown in the absence or presence of 100 µM CuSO_4_ (-Cu *vs* +Cu) identified 25 differentially expressed genes (Log_2_ Fold change ≥ 2, *p* ≤ 0.05), 17 of which were upregulated and 8 of which were downregulated in -Cu conditions (Figure 1C and Table 1). Importantly, 16 of the 17 genes upregulated in -Cu failed to display a significant change of transcript levels in the *mac1*Δ mutant. Conversely, 5 of the 8 genes downregulated in the wild-type in -Cu were also significantly downregulated in *mac1*Δ (Figure 1C).

**Table 1.** List of *F. oxysporum* genes significantly upregulated under copper limiting conditions in a Mac1-dependent manner. The closest orthologs in *S. cerevisiae* and/or *A. fumigatus* are indicated, together with a description of the protein encoded by each gene.

| Gene ID | Name | *S. cerevisiae* | *A. fumigatus* | Description |
| --- | --- | --- | --- | --- |
| Copper Transporters |  |  |  |  |
| FOXG_03101 | *ctr1a* | *ctr1* | *ctrA2* | High affinity copper transporter |
| FOXG_13082 | *ctr1b* |  |  | Putative copper transporter |
| FOXG_07770 | *ctr3* | *ctr3* | *ctrC* | High affinity copper transporter |
| Metalloreductases |  |  |  |  |
| FOXG_07769 | *fre9* |  |  | Metalloreductase |
| FOXG_18310 | *fre10* |  |  | Metalloreductase |
| FOXG_11474 | *fre11* |  |  | Metalloreductase |
| FOXG_13081 | *fre12* |  |  | Metalloreductase |
| Crm cluster |  |  |  |  |
| FOXG_09773 | *crmB* |  | *crmB* | Alcohol-O-acetyltransferase |
| FOXG_09769 | *crmC* |  | *crmC* | Putative siderophore iron transporter |
| Others |  |  |  |  |
| FOXG_04389 | *sod3* | *sod2* | *sod3* | Fe-Mn family superoxide dismutase |
| FOXG_01130 |  |  |  | Uncharacterized protein |
| FOXG_02393 |  |  |  | Uncharacterized protein |
| FOXG_06143 |  |  |  | Aminotransferase |
| FOXG_07771 |  |  |  | SAM domain-containing protein |
| FOXG_08010 |  |  |  | DJ-1_Pfpl domain-containing protein |
| FOXG_17215 |  |  |  | Acetoacetate-CoA ligase |

The 16 *Fo* genes induced under copper limitation in a Mac1-dependent manner encode known proteins involved in copper acquisition, including 3 Ctr high-affinity copper transporters (*FOXG_03101*, *FOXG_07770*, *FOXG_13082*) and 4 Fre metalloreductases (*FOXG_07769*, *FOXG_18310*, *FOXG_11474, FOXG_13081*). A BLASTp search with the predicted gene products in the genome databases of *A. fumigatus*, *Neurospora crassa*, *Pyricularia oryzae* and *S. cerevisiae* revealed that FOXG_03101 and FOXG_07770 are direct orthologs of the *S. cerevisiae* high-affinity copper transporters Ctr1 and Ctr3, respectively. Interestingly, *FOXG_13082* encodes an additional predicted Ctr transporter with high similarity to *FOXG_03101*, but a shorter protein length which was named Ctr1b (Figure S3). The three predicted Ctr copper transporters found in *Fo* are evolutionarily related with the Ctr1 and Ctr3 orthologs in other fungal species (Figure S4A). Interestingly, none of the four Fre metalloreductases upregulated in *Fo* under -Cu conditions are direct orthologs of the 8 Fre proteins reported in *S. cerevisiae* or of the FreB reductases previously annotated in *P. oryzae* and *N. crassa* (Figure S4B). We therefore named these proteins Fre9, 10, 11 and 12, with numbers ranked according to their respective transcript levels in -Cu. Among these newly annotated metalloreductases, only Fre12 is phylogenetically close to one of the previously annotated Fre proteins from *S. cerevisiae* (Fre7), whereas the others appear to be unique to filamentous fungi. Fre9 has one ortholog in *A. fumigatus* and *N. crassa* and two orthologs in *P. oryzae*, respectively, while Fre10 has a predicted ortholog in *A. fumigatus*, *N. crassa* and *P. oryzae* (Figure S4B). For Fre11 and Fre12, only one ortholog was found in *P. oryzae* and *A. fumigatus*, respectively. In addition to the high-affinity copper transporters and metalloreductases, the *Fo* genes induced under copper limitation in a Mac1-dependent manner include *sod3* encoding a Mn-dependent cytosolic superoxide dismutase, *crmB* encoding an alcohol-*O*-acetyltransferase, and *crmC* encoding a siderophore iron transporter (Figure 1C and Table 1).

To identify the Mac1 binding sites in the *Fo* genome, chromatin immunoprecipitation coupled with Next-Generation sequencing (ChIP-seq) was performed by growing the *mac1*^Stag^ strain under the same -Cu and +Cu conditions employed in the RNA-seq experiment. Using a monoclonal anti-S-tag antibody for ChIP, we identified 12 putative Mac1-binding regions in -Cu conditions whereas no Mac1-DNA interaction was detected in +Cu conditions. A comparison of these identified Mac1 binding sites defined the putative consensus DNA binding sequence of *Fo* Mac1 as 5’-DHNTGCTCANNN-3’ (D = A, G, or T; H = A, C, or T; N = any nucleotide) (Figure S5A).

The 12 Mac1-binding sites identified in the *Fo* genome map to the promoter regions of 16 genes, most of which are part of 5 gene clusters: 1) a cluster composed of the two divergently transcribed genes *ctr3*, *fre9* and the gene *FOXG_07771*, with at least one predicted Mac1-binding site in each promoter region (Figure 1D); 2) a predicted biosynthetic gene cluster composed of 5 genes: *crmA* encoding an isocyanide synthase-non ribosomal peptide synthase (ICS-NRPS)-like enzyme, *crmB*, *crmC*, *FOXG_09770* encoding a transferase family protein and *FOXG_09772* encoding a hydrolase, showing up to three putative Mac1-binding sites in each promoter except for the shared promoter of the divergently transcribed genes *FOXG_09772* and *crmB* (Figure 1E); 3) and 4) two clusters, each composed of two divergently transcribed genes sharing a common promoter region with multiple Mac1-binding sites (*fre10*/*FOXG_02393* and *fre12*/*ctr1b*) (Figure S5B and C); and 5) a gene cluster composed of *FOXG_18820* and *sod3*, which are transcribed in the same direction, containing a single Mac1-binding site between the two genes located on the opposite DNA strand (Figure S5D). Additionally, ChIP-seq identified 3 unclustered genes that are significantly upregulated in -Cu: *ctr1a*, *FOXG_01130*, and *fre11*, all of which contain Mac1-binding sites in their promoter regions (Figure S5E-G). Importantly, all the genes identified by ChIP-seq to be bound by Mac1 were also found by RNA-seq to be upregulated during -Cu, including *FOXG_09770*, *crmA*, *FOXG_09772* and *FOXG_18820* which were excluded from the main list due to their high variability between the experimental repeats (Figure 1C and Table 1).

### Mac1 is highly stable and localizes to the nucleus independently of copper status

In *S. cerevisiae*, Mac1 is rapidly degraded upon a shift from copper limitation to sufficiency(Zhu et al., 1998). To follow *mac1* expression and Mac1 protein stability in *Fo*, the *mac1*^Stag^ strain was transferred from -Cu to +Cu conditions and total RNA and protein extracts were subjected to real-time RT-qPCR and to immunoblotting with a monoclonal anti-S-tag antibody, respectively. While transcript levels were approximately halfway down 1 h after the shift, Mac1 protein levels remained unchanged even 3 h after copper addition (Figure 2A and B). Furthermore, protein levels in -Cu conditions remained stable after addition of the translation inhibitor cycloheximide (chx)(Schneider-Poetsch et al., 2010) during the entire duration of the experiment, suggesting that Mac1 turnover in *Fo* is much lower than in *S. cerevisiae*(Zhu et al., 1998) (Figure 2C). Collectively, these results suggest that *Fo* Mac1 is a highly stable protein whose activity is not primarily controlled by protein degradation.

**Figure 2.**
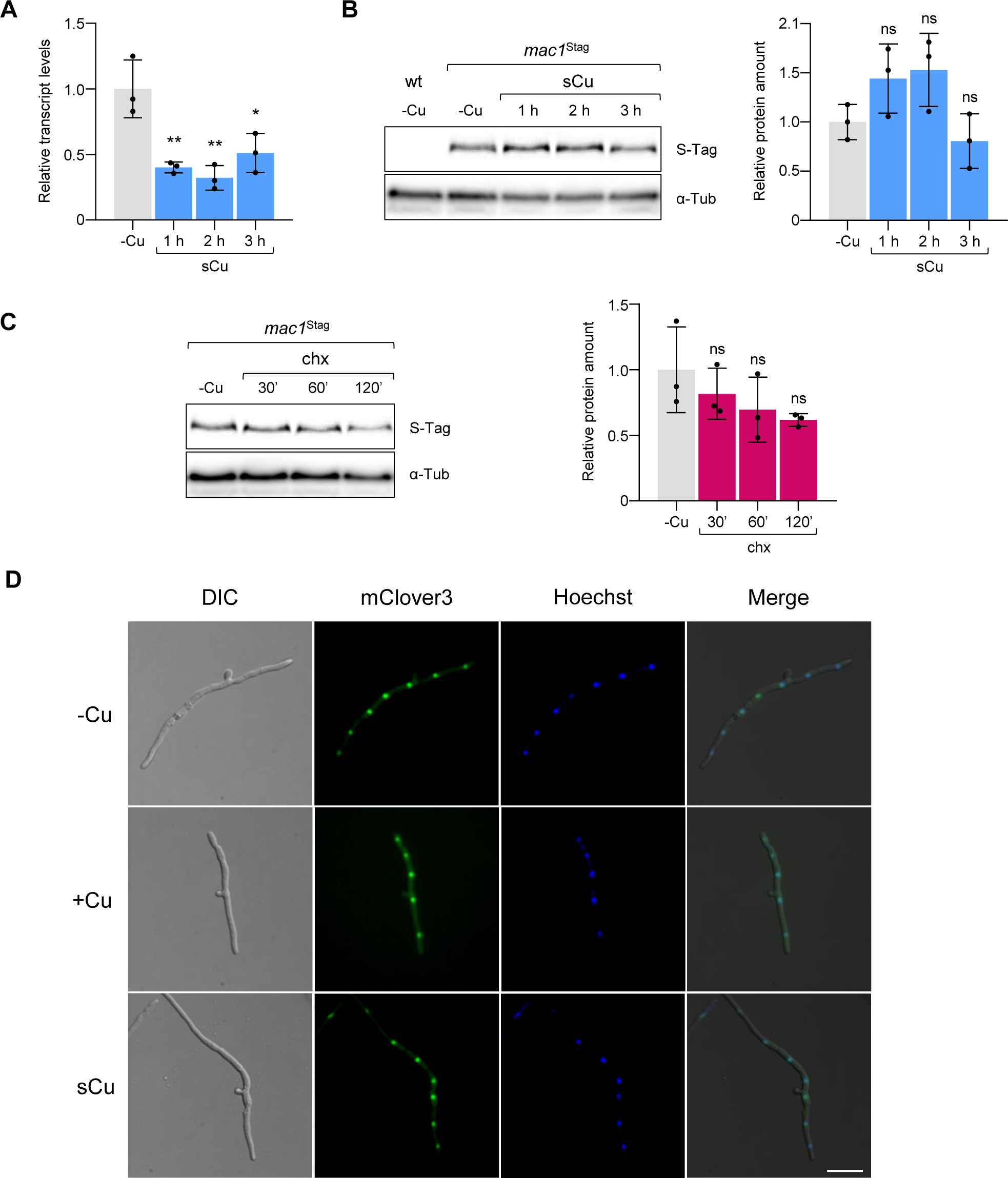
*F. oxysporum* Mac1 is highly stable and localizes to the nucleus independently of copper availability. **A.** Transcript levels of *mac1* in the *mac1*^Stag^ strain were measured by real-time RT-qPCR before and at the indicated time points after adding 20 µM CuSO_4_ (sCu) to a culture in MM-TE^-Cu^ (-Cu) medium and expressed relative to those in -Cu. Bars represent standard deviations (*n* = 3, biological replicates). *p*-values: *<0.05, **<0.01 versus -Cu according to two-tailed unpaired Student’s *t* test. **B, C.** Protein samples obtained from the *mac1*^Stag^ strain grown as described in (A) were subjected to SDS-PAGE and immunoblot analysis with anti-S-tag antibody (S-tag). Anti-α-Tubulin (α-Tub) was used as loading control. Left panels: Representative immunoblot showing Mac1 protein levels. Right panels: Densitometric quantification of Mac1 protein levels normalized to those of α-Tubulin and expressed relative to the -Cu condition. In (C), -Cu cultures were supplemented with 50 µg/ml cycloheximide (chx), instead CuSO_4_, to determine Mac1 turnover rate. Bars represent standard deviations (*n* = 3, biological replicates). *p*-values: ns>0.05 versus -Cu according to two-tailed unpaired Student’s *t* test. **D.** Subcellular localization of Mac1^clover^ was monitored after germinating the *mac1*^clover^ strain for 16 h at 28 °C in MM-TE^-Cu^ supplemented either with 100 µM (+Cu) or 2 µM CuSO_4_ (-Cu). For the copper shift experiment (sCu), 20 µM CuSO_4_ was added to the -Cu samples 10 min before imaging. Fungal nuclei were stained with Hoechst 33342. Hyphae were imaged using differential interference contrast (DIC), green fluorescence (mClover3) or blue fluorescence filters (Hoechst). The three images were merged using ImageJ v1.8. Scale bar, 25 mm.

In *A. fumigatus* and *S. pombe*, Mac1 was reported to be translocated outside of the nucleus under +Cu conditions(Beaudoin and Labbe, 2006; Park et al., 2018). To follow the subcellular localization of Mac1 in *Fo*, the *mac1*Δ mutant was transformed with a DNA construct carrying the *mac1* gene with a C-terminal fusion to the green fluorophore mClover3, driven by the constitutive *Aspergillus nidulans gpdA* promoter (Figure S6A and B). Colony growth phenotypes in -Cu and transcriptional induction of the *ctr3* and *fre9* genes of the *mac1*^clover^ strain were similar to those of the wild-type suggesting that the fluorescent tag does not interfere with Mac1 function (Figure S6C and D). Next, we performed fluorescence microscopy studies with the *mac1*^clover^ strain germinated either in - Cu or +Cu conditions or submitted to a shift from -Cu to +Cu (sCu). In contrast to previous reports in *A. fumigatus* and *S. pombe*, *Fo* Mac1^clover^ colocalized with the nuclear stain Hoechst 33342 indicating that *Fo* Mac1 is continuously present in the nucleus independent of the copper status (Figure 2D).

### Copper limitation response genes are upregulated during plant infection in a Mac1-dependent manner

To study the role of Mac1 in regulation of copper response genes during plant infection, we performed RNA-seq of tomato roots inoculated with the wild-type strain or the *mac1*Δ mutant, either at 2 or 6 days post inoculation (dpi). We first compared transcript levels of the wild-type strain during root infection to those in axenic culture under +Cu conditions and found that several copper limitation response genes were significantly upregulated during tomato plant infection (Figures 3A and S7A). These include *FOXG_17215* and *fre11* which were upregulated at 2 dpi, *FOXG_02393* which was upregulated at 6 dpi, and *fre10* and *ctr1a* which were upregulated at both infection time points. We also identified three genes that were significantly downregulated *in planta*, *crmB*, *FOXG_07771* and *FOXG_06143*. Importantly, none of the genes induced *in planta* in the wild-type were upregulated in the *mac1*Δ mutant except *FOXG_17215* (Figure 3B). Furthermore, 6 additional -Cu response genes were downregulated in the *mac1*Δ mutant compared to the wild-type *in planta*: *ctr1b*, *ctr3*, *FOXG_01130*, *FOXG_07771*, *FOXG_08010* and *fre9* (Figure 3B). Taken together, these results suggest that *Fo* faces copper limitation conditions in tomato roots and establish a key role of Mac1 in transcriptional activation of copper deficiency response genes during plant infection.

**Figure 3.**
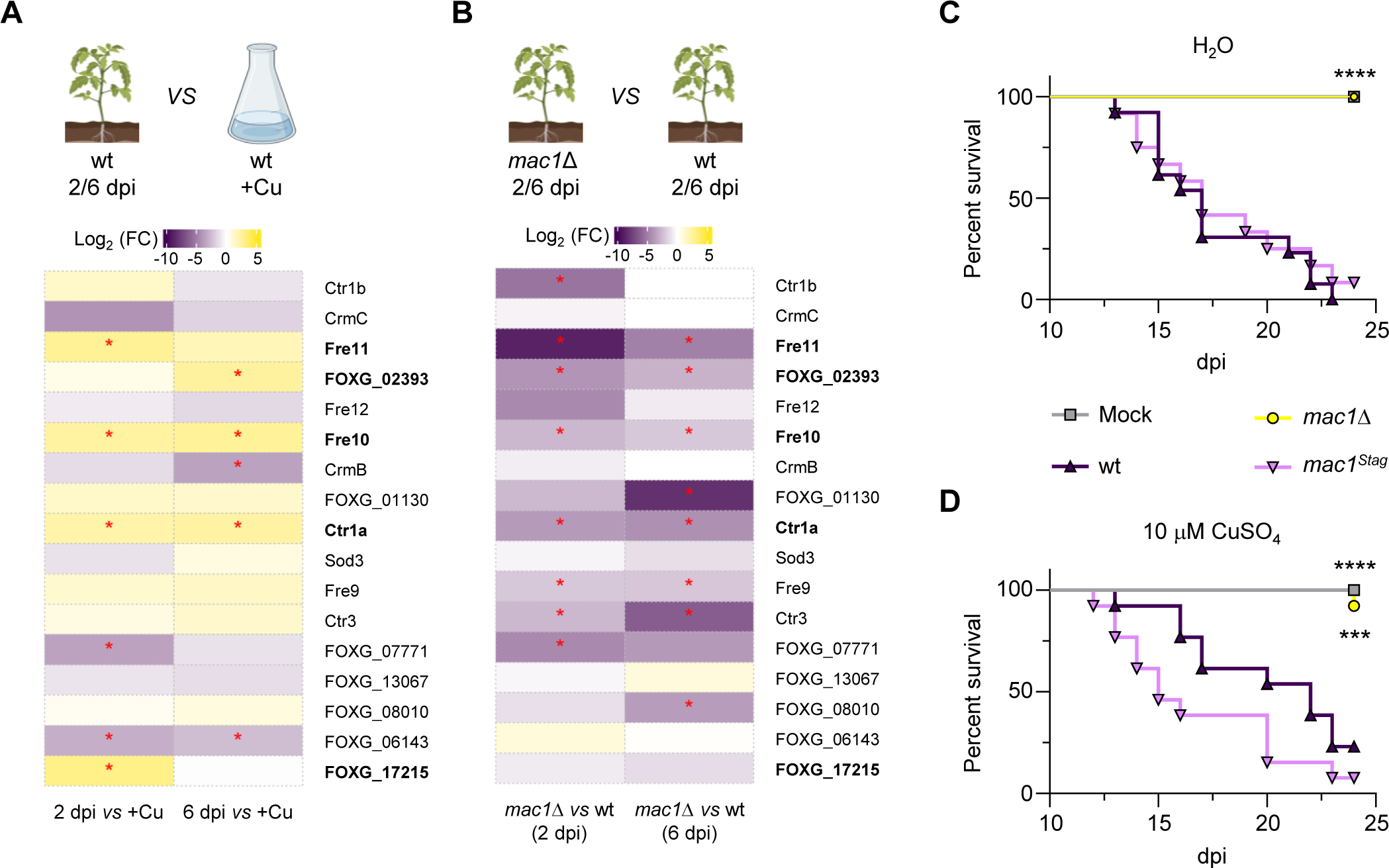
Mac1 is essential for *in planta* upregulation of copper deficiency response genes and for pathogenicity on tomato plants. **A, B.** Fold change (FC) of transcript levels of the indicated genes during infection of tomato roots by the wt at 2 (left column) or 6 (right column) days post inoculation (dpi) versus wt in axenic culture under +Cu conditions (A); or during infection of tomato roots by the *mac1*Δ strain at 2 (left column) or 6 (right column) dpi versus the wt at the same time points (B) was measured by RNA-seq. Differentially expressed genes are in the same order as in Fig. 1C. Genes upregulated both under copper limitation and plant infection conditions are in bold. *p*-value: *≤0.05 within each comparison. Data were calculated from three independent biological replicates. **C, D.** Kaplan-Meier plot showing survival of groups of 10 tomato plants (cv. Moneymaker) inoculated by dipping the roots into a suspension of 5x10^6^ microconidia/ml of the indicated fungal strain or water (Mock), planted in minipots and irrigated either with water (C) or with a 10 µM CuSO_4_ solution (D). Data shown are from one representative experiment. Experiments were performed at least three times with similar results. *p*-values: ***<0.001, ****<0.0001 versus the wt according to Log-rank (Mantel-Cox) test.

### Mac1 is essential for virulence of *Fusarium oxysporum* on tomato plants

To determine the role of *Fo* Mac1 during plant infection, roots of tomato plants were dip-inoculated with microconidia of the different fungal strains, planted in minipots, and supplied either with water or with a solution of 10 µM CuSO_4_. Under both irrigation regimes, the plants inoculated with the wild-type or the complemented strain showed mortality rates close to 100% at 25 dpi whereas those inoculated with the *mac1*Δ mutant showed no visible disease symptoms (Figure 3C and D). Using a plate invasion assay(Prados Rosales and Di Pietro, 2008), we found that the *mac1*Δ mutant was still able to penetrate across a cellophane membrane (Figure S7B). We conclude that Mac1 plays a key role in pathogenicity of *Fo* on tomato plants which is independent of invasive growth and cannot be rescued by exogenous application of copper to the roots.

### Overexpression of a copper transporter and a metalloreductase in the *mac1*Δ mutant rescues growth under copper limitation and virulence on tomato plants

We hypothesized that the essential role of Mac1 during plant infection could be related to the acquisition of copper during growth of *Fo* in the tomato root cortex and xylem. To test this idea, we first generated single and double deletion mutants in the high-affinity copper transporters Ctr1a and/or Ctr3 (Figure S8A-E). In *S. cerevisiae* and *Aspergillus*, Ctr1 and Ctr3 are functionally redundant but double deletion mutants have a severe growth phenotype in -Cu conditions(Pena et al., 2000; Park et al., 2014; Cai et al., 2017; Cai et al., 2019). Here we found that growth of the *Fo ctr3*Δ and *ctr1a*Δ single mutants and of the *ctr3*Δ*ctr1a*Δ double mutant under -Cu conditions was indistinguishable from that of the wild-type strain (Figure S8F). In line with this result, no reduction in virulence on tomato plants was observed in the single or the double mutants (Figure S8G). These results suggest the occurrence of functional redundancy in the high-affinity Ctr copper transporters of *Fo*, possibly due to the presence of the additional Ctr1 paralog Ctr1b (see Figures S3 and S4).

We next asked whether Mac1-independent expression of copper deficiency response genes in the *mac1*Δ mutant could rescue growth under Cu-conditions. This strategy is based on previous work in *Aspergillus*, where overexpression of the high-affinity copper transporters CtrA2 or CtrC (orthologs of *S. cerevisiae* Ctr1 and Ctr3, respectively) in a *mac1*Δ background partially restored growth during copper limitation(Cai et al., 2017; Cai et al., 2019). To test this idea, we generated transformants in the *mac1*Δ mutant background overexpressing either the copper transporter *ctr3* alone (*mac1*Δ*ctr3*^OE^) or both *ctr3* and the metalloreductase *fre9* (*mac1*Δ*ctr3*^OE^*fre9*^OE^) (Figures 4A and S9A-C). The *ctr3* and *fre9* genes were chosen for co-expression because they are divergently transcribed from a promoter containing 3 predicted Mac1 binding sites and are both upregulated in -Cu conditions and during plant infection in a Mac1-dependent manner (see Figure 1C and D and Figure 3A and B). Successful overexpression of *ctr3* and *fre9* in the *mac1*Δ background was confirmed in two independent *mac1*Δ*ctr3*^OE^ and *mac1*Δ*ctr3*^OE^*fre9*^OE^ transformants, respectively. RT-qPCR showed that transformants *mac1*Δ*ctr3*^OE^ #4 and *mac1*Δ*ctr3*^OE^*fre9*^OE^ #2 exhibited markedly increased *ctr3* transcript levels in -Cu conditions compared to *mac1*Δ, which were similar or even higher than those of the wild-type strain (Figure S9D). Furthermore, *fre9* transcript levels in the *mac1*Δ*ctr3*^OE^*fre9*^OE^ #2 and #4 transformants were similar to those of the wild-type strain and significantly higher than those of the *mac1*Δ mutant (Figure S9E). We next tested growth of these overexpressing strains on plates containing different levels of copper. While overexpression of *ctr3* alone was not sufficient to rescue growth of the *mac1*Δ mutant under -Cu conditions, simultaneous overexpression of *ctr3* and *fre9* restored growth in -Cu close to wild-type levels, particularly in the *mac1*Δ*ctr3*^OE^*fre9*^OE^ #2 transformant (Figure 4B). This result strongly suggest that reduction of Cu^2+^ to Cu^+^ by plasma membrane metalloreductases such as Fre9 is essential for efficient copper acquisition by *Fo*. In line with this, root infection assays revealed a direct correlation between the growth phenotype of the different strains in -Cu conditions and their ability to cause mortality on tomato plants. While the *mac1*Δ*ctr3*^OE^ transformants were only slightly more virulent than the *mac1*Δ mutant, the *mac1*Δ*ctr3*^OE^*fre9*^OE^ strains were as virulent as the wild-type (Figure 4C). Furthermore, fungal biomass at 10 dpi in stems of tomato plants inoculated with the *mac1*Δ*ctr3*^OE^*fre9*^OE^ #4 strain was not significantly different from that of the wild-type or the complemented *mac1*^Stag^ strain and significantly higher compared to plants inoculated with the *mac1*Δ mutant (Figure 4D). Similar to the wild-type strain, at 4 dpi the *mac1*Δ*ctr3*^OE^*fre9*^OE^ #4 strain labelled with the green protein mClover3 (Figure S10) was abundantly growing in the root cortex and reached the vascular bundles of the xylem, in contrast to the *mac1*Δ mutant whose growth was more sparse and remained restricted to the outermost layers of the root cortex (Figures 4E and S11). Collectively, these findings confirm that Mac1-dependent reduction and uptake of extracellular copper is essential for the progression of *Fo* from the root cortex to the vascular system and for virulence on tomato plants and demonstrate that the loss-of-virulence phenotype of the *mac1*Δ mutant is strictly associated with the inability to acquire copper.

**Figure 4.**
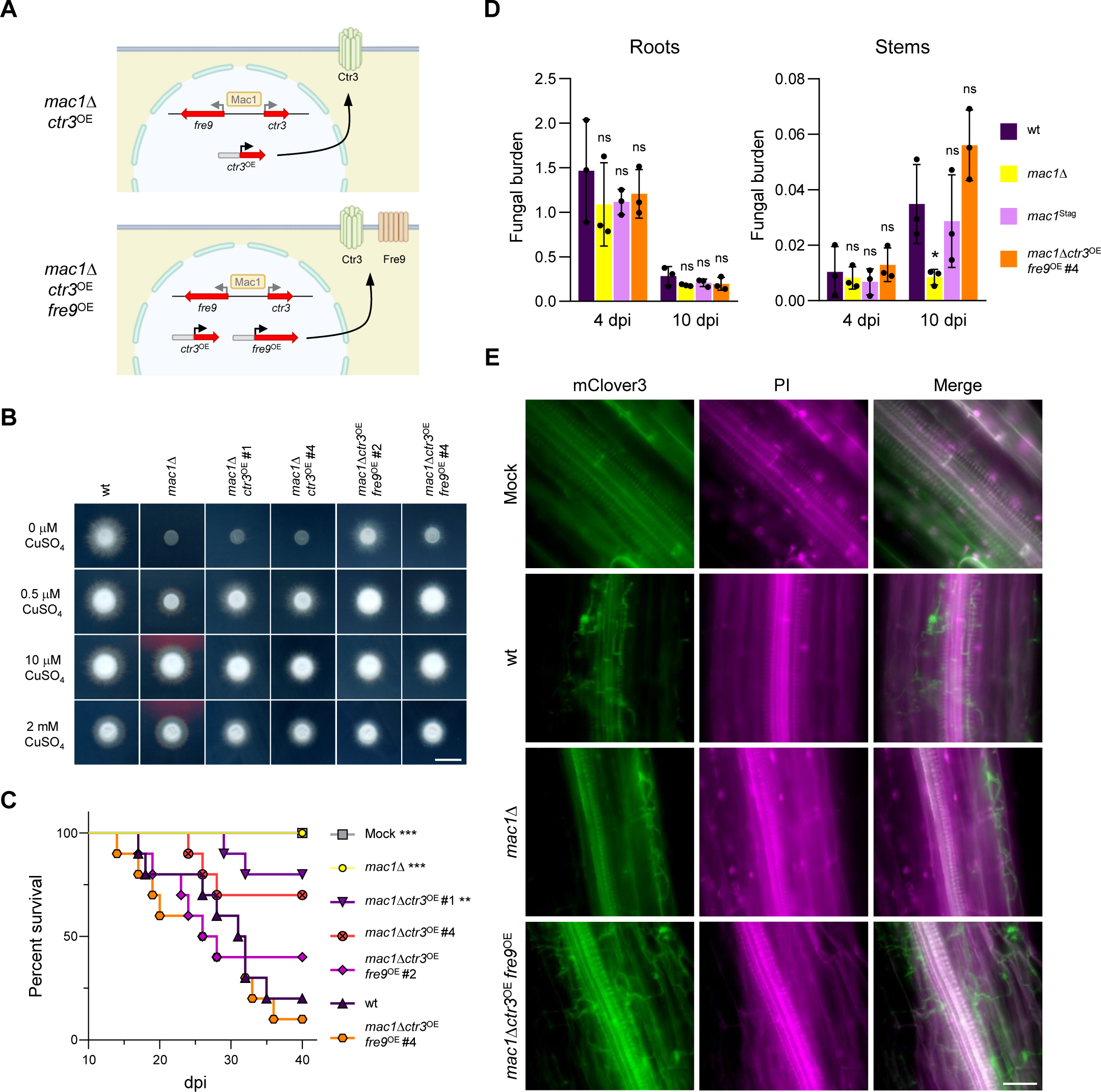
Overexpression of the copper transporter *ctr3* and the metalloreductase *fre9* in the *mac1*Δ mutant rescues growth under copper limitation and plant pathogenicity. **A.** Schematic representation of the strategy used for overexpression of the high affinity copper transporter *ctr3* alone or in combination with the metalloreductase *fre9* in the *mac1*Δ mutant driven by the constitutive *Aspergillus nidulans gpdA* promoter. **B.** Colony phenotypes of the indicated strains after 2 d growth on MM+TE^-Cu^ supplemented with the indicated concentrations of CuSO_4_. The colonies of the wt and *mac1*Δ are the same as those shown in Fig. 1A and are repeated here for clarity. Scale bar, 1 cm. **C.** Kaplan-Meier plot showing survival of groups of 10 tomato plants (cv. Momotaro) inoculated by dipping the roots into a suspension of 5x10^6^ microconidia/ml of the indicated fungal strain or water (Mock). Data shown are from one representative experiment. Experiments were performed at least two times with similar results. *p*-values: ***<0.001, ****<0.0001 versus the wt according to Log-rank (Mantel-Cox) test. **D.** Fungal burden in tomato plants inoculated with the indicated strains was measured by qPCR of the *Fol4287*-specific *six1* gene using total DNA extracted from roots (left panel) or stems (right panel) of plants at 4 or 10 dpi. Fungal burden was calculated using the 2^-ΔΔCt^ method and normalized to the tomato *gapdh* gene. Bars represent standard deviations (*n* = 3, biological replicates). *p*-value: ns>0.05, *<0.05 versus the wt in the same condition according to two-tailed unpaired Student’s *t* test. **E.** Micrographs showing fungal colonization of tomato roots (cv. Momotaro) dip-inoculated with the indicated *F. oxysporum* strains expressing 3X*Fo*-*mClover3* or water (Mock) at 4 dpi. Fungal fluorescence (mClover3, green) is overlaid with propidium iodide staining of the plant cell wall (PI, magenta). The two images were merged using ImageJ v1.8. The images shown are representative of at least three lateral secondary roots from six different tomato plants. Scale bar, 50 µm.

### Mac1-mediated copper uptake is required for virulence of *F. oxysporum* on an animal host

In the human fungal pathogens *A. fumigatus*, *C. albicans* or *C. neoformans*, loss of Mac1 leads to decreased virulence on animal hosts(Waterman et al., 2007; Cai et al., 2017; Culbertson et al., 2020). Because *Fo* can also cause disseminated infections in humans(Nucci and Anaissie, 2007), we tested the virulence phenotype of the *mac1*Δ mutant on the wax moth *Galleria mellonella*, an invertebrate model that is widely used to study microbial pathogens of humans including *Fol4287*(Navarro-Velasco et al., 2011; Navarro-Velasco et al., 2023). All larvae inoculated with the wild-type or the *mac1*^Stag^ strain were dead at 2 dpi while most larvae inoculated with *mac1*Δ remained alive at 5 dpi (Figure S12A). Importantly, the ability to cause mortality on *G. mellonella* was rescued to wild-type levels in the two *mac1*Δ*ctr3*^OE^*fre9*^OE^ transformants (Figure S12B), demonstrating that Mac1-dependent copper acquisition is required for full virulence of *F. oxysporum* on this animal host.

## DISCUSSION

The copper-sensing transcription factor Mac1 was identified more than thirty years ago in *S. cerevisiae*(Jungmann et al., 1993), in which copper homeostasis has been studied in detail. Mac1 activates the transcription of genes encoding metalloreductases and high-affinity copper transporters which are required for efficient copper acquisition under copper limitation, and its loss provokes a severe phenotype in -Cu conditions(Smith et al., 2017). While a role of Mac1 in fungal virulence on animal hosts has been demonstrated(Waterman et al., 2007; Sun et al., 2014; Cai et al., 2017; Raffa et al., 2019; Culbertson et al., 2020; Ray and Rappleye, 2022); its role during plant infection has not been studied before.

Here we functionally characterized Mac1 in *Fo*, an important fungal phytopathogen. The structural organization of *Fo* Mac1 is similar to its orthologs in other fungi, with an N-terminal copper fist DNA-binding domain containing the characteristic C, RGHR and GRP residues, and two C-terminal Cys-rich motifs. Furthermore, the inability of the *Fo mac1*Δ mutant to grow in -Cu conditions confirms that the function of Mac1 is highly conserved across different fungi, which is in line with a previous study showing that the *mac1* gene from *A. fumigatus* can functionally complement a *mac1*Δ mutant of *S. cerevisiae*(Park et al., 2017). By contrast, we found that the post-translational regulation of Mac1 activity in *Fo* differs from that reported in other fungi where inactivation under copper sufficiency occurs mainly by protein degradation and/or cytoplasmic retention, analogous to what has been reported for the iron-responsive transcription factor HapX(Lopez-Berges et al., 2021). In *S. cerevisiae*, Mac1 is stable at low copper concentrations but rapidly degraded at concentrations above 10 µM CuSO_4(Zhu et al., 1998)_. Furthermore, Mac1 in *A. fumigatus* and *S. pombe* localizes to the nucleus in -Cu but is present in the cytoplasm during copper sufficiency(Beaudoin and Labbe, 2006; Park et al., 2018). Here we found that *Fo* Mac1 has a low turnover rate, is highly stable and remains predominantly localized in the nucleus during +Cu, suggesting that its activity is not regulated by protein degradation or cytoplasmic retention. The most likely explanation for the high Mac1 DNA binding specificity under -Cu conditions is a mechanism previously proposed in *S. pombe*(Beaudoin and Labbe, 2006), where copper induces a conformational change in the protein that promotes physical interaction between the N-terminal DNA-binding domain and a Cys-rich motif thereby preventing binding of the transcription factor to its target sites in the genome.

Using a combination of RNA-seq and ChIP-seq, we detected direct DNA binding of *Fo* Mac1 only under -Cu conditions and found that it transcriptionally activates a group of genes involved in copper acquisition (Ctr copper transporters and Fre reductases), reactive oxygen species (ROS) detoxification (Sod3), and production of isocyanine metabolites (Crm gene cluster). In other fungi, Mac1 also was shown to activate copper acquisition genes, including the orthologs of *S. cerevisiae ctr1* and *ctr3*(Smith et al., 2017; Raffa et al., 2019). In addition, *Fo* has a third high affinity copper transporter, *ctr1b*, which is also present in the rice blast fungus *P. oryzae* and is transcriptionally induced by Mac1 during -Cu conditions. Our finding that the *ctr1*/*ctr3* double mutant is still able to grow under -Cu conditions strongly suggests that Ctr1b is functionally redundant with Ctr1a and Ctr3.

In *S. cerevisiae*, the two copper reductases Fre1 and Fre7 are transcriptionally induced during -Cu in a Mac1-dependent manner(Qi et al., 2012). Interestingly, the direct *fre1* and *fre7* orthologs were not upregulated in *Fo* during -Cu. Instead, we found 4 Fre reductases that were direct targets of *Fo* Mac1, which have no close orthologs in *S. cerevisiae* and appear to be specific for filamentous fungi. Two of these genes, *fre9* and *fre12,* are clustered and divergently transcribed with the high-affinity copper transporters *ctr3* and *ctr1b*, respectively, suggesting that they are functionally related. Apart from the genes involved in copper acquisition, we found that *Fo* Mac1 directly activates *sod3* encoding a cytosolic Mn-dependent superoxide dismutase. In *C. albicans*, Sod3 was shown to be important for the maintenance of ROS homeostasis under -Cu conditions, due to the reduced activity of the most abundant cytosolic superoxide dismutase Sod1, which requires copper as cofactor(Culbertson et al., 2020). Furthermore, several genes in the *crm* biosynthetic gene cluster of *Fo* were identified as direct Mac1 targets in *Fo*. The *crm* cluster was recently shown to be induced under copper starvation in *A. fumigatus* and is conserved in a wide range of pathogenic and non-pathogenic fungi. It encodes, among others, an isocyanide synthase that contributes to two distinct biosynthetic pathways whose final products have antibacterial properties(Lim et al., 2018; Won et al., 2022). Importantly, several of the direct Mac1 targets in *Fo* were previously reported to contribute to virulence in human fungal pathogens(Raffa et al., 2019; Won et al., 2022).

Our results demonstrate for the first time an essential role of Mac1-mediated copper uptake during plant infection by a fungal pathogen. The observation that exogenous addition of 10 µM CuSO_4_, a concentration sufficient to rescue growth of the *mac1*Δ mutant on plates, failed to restore virulence on tomato plants led us to hypothesize that *Fo* Mac1 could directly or indirectly activate virulence-related genes other than those related to copper limitation. However, comparative RNA-seq analysis during tomato root infection demonstrated that the transcripts downregulated in *mac1*Δ mutant versus wild-type *in planta* all correspond to genes induced under copper starvation, most of which are directly related to copper acquisition. Together with the finding that simultaneous overexpression of the high-affinity copper transporter Ctr3 and the metalloreductase Fre9, but not of Ctr3 alone, rescues growth in -Cu conditions and virulence of the *mac1*Δ mutant, our results strongly indicate that both extracellular copper reduction as well as copper uptake are essential during these processes, a conclusion that is further supported by the direct correlation observed between the ability of the different *Fo* strains to grow in -Cu conditions and to cause wilt disease on tomato plants. Fungal burden experiments and fluorescence microscopy studies revealed that *in planta* growth of the *mac1*Δ mutant was largely restricted to the outermost layers of the cortex, in stark contrast to the wild-type or the *mac1*Δ*ctr3*^OE^*fre9*^OE^ strains who were able to progress towards the innermost root layers and reach the vascular system. Interestingly, the acquisition mechanism of copper by tomato roots has been proposed to function similar to that of fungi, whereby Cu^2+^ is initially reduced to Cu^+^ which is subsequently taken up by the root(Ryan et al., 2013). However, for loading of copper into the xylem re-oxidation of Cu^+^ to Cu^2+^ is required, followed by complexation with the mugineic acid-derived metal chelate nicotianamine for further translocation(Ryan et al., 2013). This redox-selective process implies that Cu^+^ is available in the root cortex but largely absent during later stages of fungal infection in the xylem vessels due to its oxidation to Cu^2+^, explaining the residual growth of the *mac1*Δ mutant in the outer root layers and its failure to colonize the xylem and progress to the plant stems and to cause wilt disease (Figure 5). Interestingly, we found that efficient copper reduction and uptake are also essential for virulence of *Fo* on *G. mellonella*, establishing a conserved role of copper acquisition in fungal pathogenicity on hosts from different kingdoms of life. These results suggest that targeting copper acquisition may represent a promising strategy for controlling diseases caused by phytopathogenic fungi.

**Figure 5.**
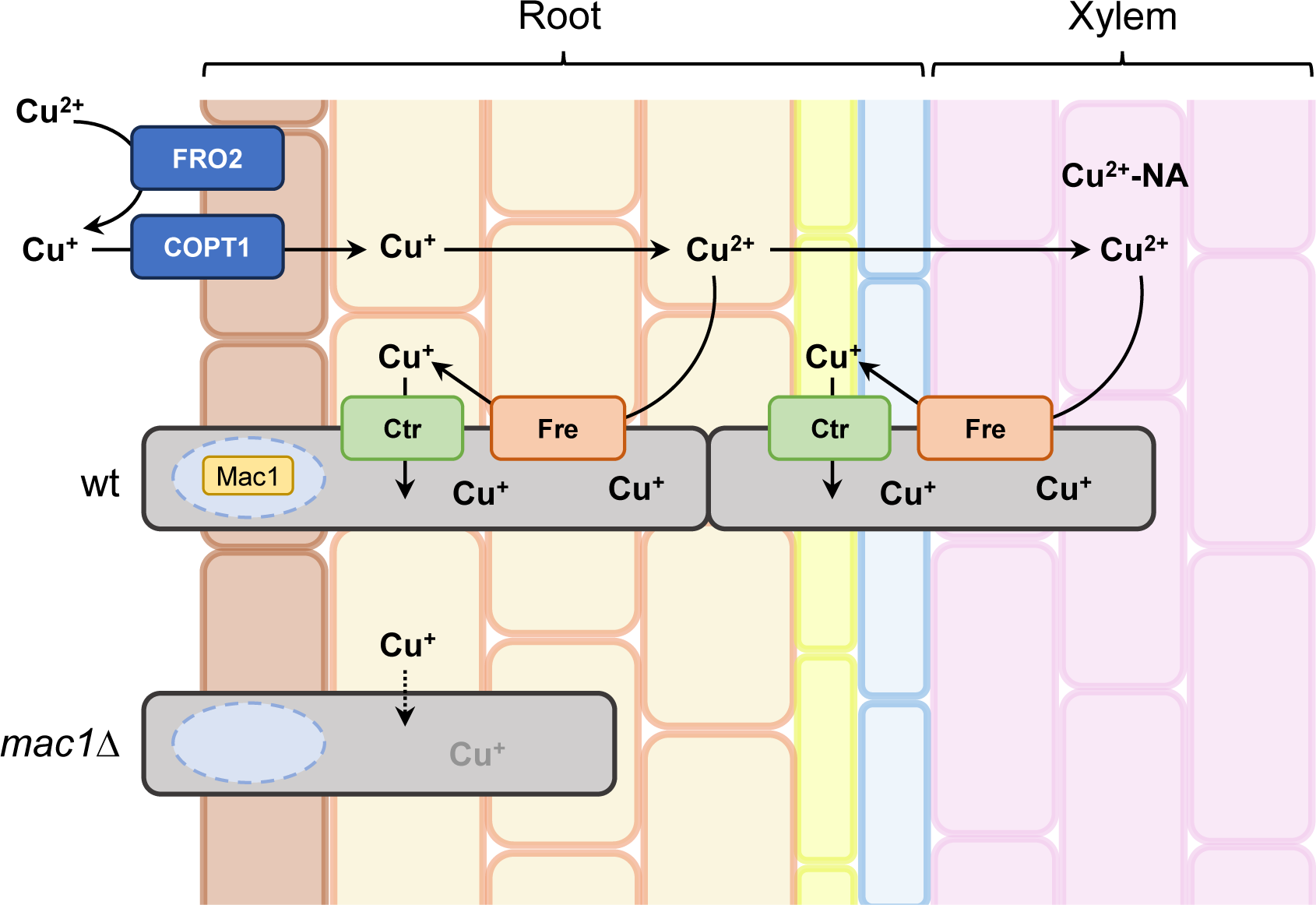
*In planta* copper reduction and uptake by *F. oxysporum* is essential for root infection and vascular colonization of tomato plants. Copper (Cu^2+^) is reduced and then taken up by tomato roots as Cu^+^. However, Cu^+^ is re-oxidized to Cu^2+^ for xylem loading and complexed with the metal chelate nicotianamine (NA) for translocation^43^. Successful root infection and vascular colonization of *F. oxysporum* therefore depends on *in planta* copper uptake, which in turn requires upregulation of copper reductase and transporter genes by direct binding of the transcription factor Mac1.

## METHODS

### *Fusarium oxysporum* strains

*Fusarium oxysporum* f. sp. *lycopersici* 4287 (*Fol4287*) was used in all experiments. All the strains for this study were generated in this genetic background and are listed in Table S1. Primers and plasmids used in this work are listed in Tables S2 and S3. Targeted deletion of the *mac1* (*FOXG_03227*), *ctr1a* (*FOXG_03101*), and *ctr3* (*FOXG_07770*) genes was performed by homologous gene replacement with the hygromycin B (*mac1* and *ctr3*) or the neomycin (*ctr1a*) resistant cassettes using the split marker method(Goswami, 2012) (Figures S2A and S9A and C). Additionally, *ctr1a* was deleted in a *ctr3*Δ background. Briefly, two PCR fragments encompassing 1.5 kb of the 5’- and 3’-flanking regions were amplified by PCR with primer pairs Mac1-5’-F + Mac1-5’-R and Mac1-3’-F + Mac1-3’-R, Ctr3-5’-F + Ctr3-5’-R and Ctr3-3’-F + Ctr3-3’-R, and Ctr1a-5’-F + Ctr1a-5’-R and Ctr1a-3’-F+ Ctr1a-3’-R, for *mac1*, *ctr3* and *ctr1a*, respectively. The amplified fragments were then fused to the hygromycin/neomycin resistance cassettes, previously amplified from pAN-7.1(Punt et al., 1987) or pGEMT-Neo(Fernandes et al., 2017) plasmids, with primer pairs Mac1-hph-F + Mac1-hph-R, Ctr3-hph-F + Ctr3-hph-R or Ctr1a-neo-F + Ctr1a-neo-R, using the fusion primer combinations Mac1-5’-Fn + HygY and Mac1-3’-Rn + HygG, Ctr3-5’-Fn + HygY and Ctr3-3’-Rn + HygG or Ctr1a-5’-Fn + NeoY, and Ctr1a-3’-Rn + NeoG. The two resulting DNA constructs for each target gene were used to co-transform freshly prepared *F. oxysporum* protoplasts. The obtained transformants were purified by two rounds of monoconidial isolation as described(Di Pietro and Roncero, 1998). Hygromycin/geneticin-resistant transformants were analyzed by Southern blot analysis with gene-specific probes (Figures S2B and S8B, D and E).

For complementation of the *mac1*Δ mutant #1, a 3,938 bp DNA fragment containing the wild-type *mac1* ORF fused at the 3’ end with a DNA sequence encoding for the S-tag oligopeptide (*mac1*^Stag^) was used (Figure S2C). To this aim, a 72 bp sequence encoding a 4X-GA linker and the oligopeptide S-tag was amplified from plasmid *Stag::pyrG*(Pinar et al., 2015) with the primer pair Mac1-Stag-F + Mac1-Stag-R. Next, two fragments containing the *mac1* ORF without the Stop codon preceded by 1,167 bp of its 5’ region, and 1,095 bp of its 3’ region were amplified from *Fol4287* genomic DNA with the primers Mac1-5’-F + Mac1-ORF-STOP-R and Mac1-TER-F + Mac1-3’-R, respectively. The three obtained DNA fragments were fused by PCR using the primer pair Mac1-5’-Fn + Mac1-3’-Rn. Taking advantage of the inability of *mac1*Δ to grow under -Cu conditions (see Figure 1A), no selection marker was used. Transformants that grew in the absence of copper (MM+TE^-Cu^) were inoculated in plates with and without hygromycin. Among all the transformants capable of growing in MM+TE^-Cu^, one had lost hygromycin resistance indicating *in locus* integration of the complementation construct. This strain (hereafter called *mac1*^Stag^) was further analyzed by PCR with locus-specific primers (Figure S2D).

The *mac1*^clover^ strain was generated by the co-transformation of *mac1*Δ protoplasts with the *Phleo*^R^ cassette, amplified from plasmid pAN8-1(Mattern et al., 1988) using the primer pair Gpda15B + TrpC8B, and the *mac1*^clover^ allele (Figure S6A). To generate this DNA construct, four DNA fragments were obtained by PCR: the *A. nidulans gpdA* promoter and the 1X*Fo-mClover3* gene were both amplified from plasmid pUC57-1X*FomClover3*(Redkar et al., 2022) with primer pairs Gpda15B + Gpda-Mac1-R and Mac1-Clover-F + Clover-Mac1-Ter-R, respectively. The *mac1* ORF without the Stop codon and a 1,124 bp fragment of the 3’ flanking region of *mac1* were amplified from *Fol4287* genomic DNA using primer pairs Mac1-ATG-F + Mac1-ORF-STOP-R and Mac1-TER-F + Mac1-3’-R, respectively. The four obtained DNA fragments were fused by PCR using the primer pair Gpda15Bnest + Mac1-3’-Rn. PCR analysis with the primers Mac1-qPCR-F and EYFPrev identified four independent transformants showing a PCR amplification product with the expected size of 1,418 bp (Figure S6B).

To achieve constitutive expression of *ctr3* and *fre9* on the *mac1*Δ background, the CDSs followed by approximately 1.3 kb of the terminator regions were amplified using primer pairs Gpda-Ctr3 + Ctr3-3’-R or Gpda-Fre9 + Fre9-3’-R, respectively. Then the *gpdA* promoter of *A. nidulans* was amplified from plasmid pAN7-1(Punt et al., 1987) with the primer pair GpdA15B + GpdA9 and fused to the 5’ ends of the *ctr3* or the *fre9* fragments using the primers Gpda15nest + Ctr3-3’-Rn or Fre9-3’-Rn, to generate the DNA constructs *ctr3*^OE^ and *fre9*^OE^, respectively (Figure S9A). Next, *mac1*Δ protoplast were co-transformed with the *Nat*^R^ cassette, amplified from plasmid pDNat(Kopke et al., 2010) with the primer pair M13-F + M13-R, together with the *ctr3*^OE^ DNA construct (for generating *mac1*Δ*ctr3*^OE^) or with both *ctr3*^OE^ and *fre9*^OE^ DNA fragments (for generating *mac1*Δ*ctr3*^OE^*fre9*^OE^). PCR analysis of two independent nourseothricin-resistant transformants from each transformation experiment, with specific the primers Gpda4 + Ctr3-FOXG_07770-R or Fre9-3’-Rn, confirmed the presence of the *ctr3*^OE^ or the *fre9*^OE^ DNA constructs in these strains (Figures S8B and C).

For fluorescence microscopy assays under infection conditions, we generated *mac1*Δ or *mac1*Δ*ctr3*^OE^*fre9*^OE^ #4 strains expressing 3 copies of *Fo-mClover3* by co-transformation with the *Phleo*^R^ cassette, amplified from plasmid pAN8-1(Mattern et al., 1988) using the primers Gpda15B + TrpC8B, and the *Fo-mClover3* expression cassette, amplified from plasmid pUC57-3X*FomClover*3(Redkar et al., 2022) with primer pair Gpda15B + SV40rev (Figure S10A). PCR analysis with the gene-specific primers Gpda4 + 3XFLAGrev revealed the presence of the *Fo-*3X*mClover3* expression cassette in four *mac1*Δ-3X*mClover* transformants and in two *mac1*Δ*ctr3*^OE^*fre9*^OE^-3X*mClover* transformants (Figures S10B and C). *mac1*Δ-3X*mClover* #11 and *mac1*Δ*ctr3*^OE^*fre9*^OE^-3X*mClover* #1 were selected for microscopy studies because they presented the highest fluorescence intensity.

### Culture conditions

Fungal strains were stored at -80 °C as microconidial suspensions in 30% glycerol (v/v). For microconidia production and DNA extraction, strains were grown for 3-4 d in liquid potato dextrose broth (PDB) at 28 °C and 170 rpm. When needed, appropriate antibiotics (hygromycin B at 20 μg/ml, geneticin at 10 μg/ml, phleomycin at 4 μg/ml, and nourseothricin at 2.5 μg/ml) were added to the culture medium.

For phenotypic analysis of colony growth, 5 μl drops of 10^7^ microconidia/ml in water were spotted onto 20 mM L-glutamine minimal medium with -Cu trace elements (MM+TE^-Cu^) solid plates supplemented with 0 to 2 mM CuSO_4_. To determine the fungal biomass production, 2.5x10^6^ microconidia/ml were germinated in liquid MM+TE^-Cu^ supplemented, or not, with 10 µM CuSO_4_ for 16 h at 28 °C and 170 rpm. The obtained mycelium was lyophilized and weighed.

Axenic liquid cultures at specific copper concentrations were grown as previously described(López-Berges, 2020). 2.5x10^6^/ml freshly obtained microconidia were inoculated in PDB and cultured for 15 h at 28 °C and 170 rpm. Germlings were filtered using a Monodur filter membrane, washed three times with sterile Milli-Q water, collected with a sterile spatula, divided in two flasks containing pH 6.5 MM+TE^-Cu^ with the desired CuSO_4_ concentration and incubated for 6 h at 28 °C and 170 rpm. In some experiments, a shift from -Cu to +Cu was carried out by the addition of CuSO_4_ after 6 h of incubation in MM+TE^-Cu^. Copper concentrations used in each experiment are indicated in the figures.

When needed, protein biosynthesis was blocked using 50 µg/ml of the translation inhibitor cycloheximide (chx) (Milipore^®^). All glassware used was pre-washed with 3.5% HCl for 30 min and rinsed five times with distilled water to remove any traces of metal adhering to the glass.

### Nucleic acid manipulation and quantitative real-time reverse transcription-polymerase chain reaction (RT-qPCR) analysis

Genomic DNA was extracted as previously reported(Torres et al., 1993). DNA was quantified in a Nanodrop^®^ ND1000 spectrophotometer at 260 nm and 280 nm wavelengths. The quality of the DNA was monitored by electrophoresis in 0.7% agarose gels (w/v). PCR amplification reactions were performed using different thermostable Taq DNA polymerases depending on the experiment and the expected fragment size. The enzymes Expand^TM^ High Fidelity PCR System (Roche) or Phusion^®^ High-Fidelity DNA Polymerase (New England Biolabs^®^) were used for reactions where high fidelity PCR amplification was required. For routine PCRs and Southern blot probes, the thermostable BioTaq^TM^ DNA Polymerase (Meridian Bioscience^®^) was used. The amplifications were performed according to the manufacturer’s instructions in a T100 Thermal Cycler (Bio-Rad).

To measure transcript levels of the desired genes, total RNA was isolated from snap frozen tissue of three biological replicates and used for reverse transcription quantitative PCR (RT-qPCR) analysis as described(Lopez-Berges et al., 2012; López-Berges, 2020). Briefly, RNA was extracted using the Tripure Reagent and treated with DNase (both from Roche). The resulting RNA was reverse transcribed with the iScript^TM^ cDNA Synthesis Kit (Bio-Rad) to synthesize the cDNA, and qPCR was carried out using the FastStart Essential DNA Green Master (Roche) in a CFX Connect Real-Time System (Bio-Rad) according to the manufactureŕs instruction. Data were analyzed using the double delta Ct method(Livak and Schmittgen, 2001; Pfaffl, 2001) by calculating the relative transcript level normalized to the *act1* gene (*FOXG_01569*).

### RNA sequencing analysis

RNA-seq analysis was carried out using RNA isolated from samples obtained from axenic cultures (supplemented or not with 100 µM CuSO_4_) and from infected tomato roots. For this, roots of 2-week-old tomato plants were inoculated with the indicated *F. oxysporum* strains and harvested either at 2 or 6 days post inoculation (dpi). Groups of three roots were sampled together and considered as one biological replicate. Once collected, samples (either mycelium or tomato roots) were frozen in liquid nitrogen and lyophilized. RNA was extracted using the RNeasy^®^ Plant Mini Kit (Qiagen) as described(Redkar et al., 2022) and treated with DNase I using the Turbo DNA-Free^TM^ Kit (Invitrogen^TM^) according to the manufacturer’s instructions. RNA sequencing was performed by Novogene, UK. For library preparation mRNA was captured through poly-A enrichment on the total RNA, and a TruSeq RNA Library Preparation Kit (Illumina, USA) was used to build the libraries according to the manufacturer’s protocol. Libraries were sequenced on a NovaSeq6000 sequencing platform (Illumina). Paired-end 150 bp reads were obtained for each RNA-seq library. Raw reads were produced from the original image data by base calling. Reads containing adaptors, highly ‘N’ containing reads (>10% of unknown bases) and low-quality reads (more than 50% bases with quality value of <5%) were removed. After data filtering, on average, ∼99.3% clean reads remained. Transcript quantification was performed with Salmon v1.6.0(Patro et al., 2017). RNA-seq paired-end read data sets were quasimapped against the reference transcriptome of *Fusarium oxysporum* f. sp. *lycopersici 4287* (GCF_000149955.1_ASM14995v2_rna.fna, obtained from NCBI RefSeq). Raw gene counts were used to evaluate the level of correlation between biological replicates using Pearson’s correlation matrix with corrplot R package v0.92(Wei and Simko, 2021) (Figure S13).

Differential gene expression analysis at transcript level was analyzed using DESeq2 R package v1.40.2(Love et al., 2014). Transformed raw counts (vst function) were used for Principal Component Analysis (PCA) using prcomp function from stats R package(Team, 2023) (Figure S14). A cut-off of absolute FC [log2 fold change] ≥ 2 and adjusted p-value ≤ 0.05 by Benjamini and Hochberg method were used to identify differentially expressed genes (DEGs). Genes with less than 1 transcript per million (TPM) across all samples were considered lowly expressed and ignored in the analysis. Intersection between DEGs in the wild-type strain in -Cu and root samples at 2 and 6 dpi, against +Cu as control were calculated and visualized with ComplexUpset R package v1.3.3(Lex et al., 2014; Krassowski, 2020). Genes were hierarchically clustered based on FC using the Heatmap function from the ComplexHeatmap package R package v2.16.0(Gu et al., 2016). The R statistical language and environment v4.3.0 was used for RNA-seq data analysis and visualization(Team, 2023). Scripts used are available at https://github.com/mvapontes/palosfernandez_et_al_plant_2023.

### Chromatin immunoprecipitation-coupled sequencing analysis (ChIP)

ChIP-seq analysis of fungal cells grown in submerged liquid cultures followed the procedures described(Gacek-Matthews et al., 2016). Briefly, DNA was crosslinked to proteins by adding 1% formaldehyde (v/v) to the axenic cultures and incubating them for 15 min at 28 °C and 170 rpm. Crosslinking was stopped by the addition of 125 mM glycine (final concentration) and 5 min incubation with shaking. The mycelium was collected by filtration through a Monodur nylon filter and flash-frozen in liquid nitrogen. Mycelium was ground in liquid nitrogen with a mortar and a pestle. ChIP was carried out as described(Reyes-Dominguez et al., 2012) with minor modifications. The monoclonal anti-S-tag antibody (SAB2702227, Sigma-Aldrich^®^) was used. Precipitation of the protein-antibody conjugate was performed with Dynabeads^TM^ Protein G (10003D, Thermo Fisher Scientific™). Chromatin-bead complexes were washed three times with Low-salt buffer followed by one wash with 1 ml High-salt buffer and eluted in TES buffer (50 mM TRIS-HCl pH 8, 10 mM EDTA pH 8, 1% SDS). Chromatin was treated with Proteinase K (MBI) and DNA purification was done using the PCR and DNA Cleanup Kit (Monarch^®^). All experiments were performed in biological triplicates.

The obtained DNA was sent for sequencing at the Vienna BioCenter Core Facilities (Vienna, Austria). Paired-end sequencing was performed using a NextSeq550 PE75 Illumina sequencer. Obtained sequences were de-multiplexed, quality controlled, filtered using trimmomatic 0.36(Bolger et al., 2014) and mapped on the already available *Fusarium oxysporum* f. sp. *lycopersici* 4287 genome assembly (GCF_000149955.1_ASM14995v2_genomic.fna from NCBI RefSeq). Mapping was performed using BWA(Li and Durbin, 2009) and further processing was done using samtools 1.7 and bedtools v2.27.1 to obtain normalized genome coverage tracks.

For identification of Mac1 binding site coverage, tracks were loaded into R and peaks were identified using R function locate_peak_height (https://github.com/symbiocyte/MNase). Peaks identified in -Cu and +Cu were visually selected using IGB v9.1.10 (https://www.bioviz.org/) resulting in 12 Mac1 specific peaks. Genomic sequences around these peak locations (500 bp) were exported and submitted to NCBI-BLAST homology searches against nt database for the identification of conserved regions. Within these regions the consensus sequence TGCTCA could be identified. A search of the motif was conducted using FIMO(Grant et al., 2011) online tool at MEM suite with default parameters.

ChIP-seq and RNA-seq coverage plots of Mac1 selected genomic regions were created using kpPlotBAMCoverage function from KaryoplotR(Gel and Serra, 2017) Bioconductor package version 1.26.0 in R statistical language(Team, 2023). The employed script is available at https://github.com/mvapontes/palosfernandez_et_al_plant_2023.

### Sequence search and phylogenetic analysis

In silico gene and protein searches of *Fol4287* and related fungal species was performed using the BLAST algorithm(Altschul et al., 1990) from the National Center for Biotechnology Information (NCBI; http://www.ncbi.nlm.nih.gov), Fungal and Oomycete genome database (FungiDB; https://fungidb.org/fungidb/app) and Saccharomyces Genome Database (SGD; https://www.yeastgenome.org). Protein domain prediction was done using the Prosite database (ExPASy; https://prosite.expasy.org), Pfam (http://pfam.xfam.org), InterPro (https://www.ebi.ac.uk/interpro/) and NCBI Conserved Domain Search (https://www.ncbi.nlm.nih.gov/Structure/cdd/wrpsb.cgi). Protein alignments were done using the BioEdit software v7.7.1.

For phylogenetic analysis, genome mining of *Fol4287* against selected proteins was performed using BLASTp. Results were manually curated based on percentage of identity and e-value. MAFFT v7.453(Katoh and Standley, 2013) with default parameters was used to align protein sequences. Manually curated alignments were used to generate phylogenetic trees using MEGA v11(Tamura et al., 2021) with Maximum Likelihood method and JTT matrix-based model. The bootstrap consensus phylogenetic tree was inferred from 1000 replicates.

### Fluorescence microscopy

For studying the subcellular localization of Mac1, 2.5x10^6^ freshly obtained microconidia/ml of the *mac1*^clover^ strain were germinated for 16 h at 28 °C and 170 rpm in 20 mM L-glutamine MM+TE^-Cu^ with 100 µM CuSO_4_ (+Cu) or 2 µM CuSO_4_ (-Cu). In some experiments, -Cu cultures were shifted to +Cu (20 µM CuSO_4_) during 10 min before imaging. Fungal nuclei were stained 5 min before imaging with 2 µg/ml Hoechst 33342 (Invitrogen™) in water in the dark. Wide-field fluorescence imaging was performed with a Zeiss Axio Imager M2 microscope equipped with a Photometrics Evolve EMCCD camera, using the 40X oil objective. Fo-mClover and Hoechst 33342 fluorescence were visualized at an excitation of 459 and 352 nm, and emission detected at 519 and 461 nm, respectively.

For microscopic observations of *F. oxysporum* during tomato plant infection, roots of 2-week-old tomato seedlings inoculated with the *F. oxysporum* strains of interest were collected at 4 dpi and secondary lateral roots were sampled. To visualize plant cell walls, samples were stained with 2 mg/ml propidium iodide (PI) (Sigma-Aldrich^®^) in water in the dark for 15 min before imaging. Wide-field fluorescence imaging was performed with a Zeiss Axio Imager M2 microscope equipped with a Photometrics Evolve EMCCD camera, using the 40X oil objective. Fo-mClover and PI fluorescence were visualized at an excitation of 459 and 587 nm, and emission detected at 519 and 610 nm, respectively.

### Western blot analysis

Proteins were extracted using a reported procedure(Hervas-Aguilar and Penalva, 2010; Lopez-Berges et al., 2016) involving solubilization from lyophilized mycelial biomass with NaOH, followed by precipitation with trichloroacetic acid (TCA). Aliquots were resolved in 10% SDS-polyacrylamide gels (Bio-Rad) and transferred to nitrocellulose membranes with a Trans-Blot^®^ Turbo^TM^ Transfer System (Bio-Rad) for blotting. Western blots were reacted with monoclonal anti-S-tag (1:5,000; SAB2702227, Sigma-Aldrich^®^) as primary antibody and with polyclonal anti-mouse IgG peroxidase (1:5,000; #7076, Cell Signalling Technology^®^) as secondary antibody. Tubulin, used as loading control, was detected with monoclonal anti-α-Tub (1:5,000; T9026, Sigma-Aldrich^®^) as primary antibody and with polyclonal anti-mouse IgG peroxidase (1:5,000; #7076, Cell Signalling Technology^®^) as secondary antibody. Proteins were detected by chemiluminescence using ECL Select^TM^ Western blotting Detection reagent (GE Healthcare, Amersham^TM^) and a Fujifilm LAS-3000 camera.

### Cellophane penetration assay

The cellophane penetration assay was performed as previously described(Prados Rosales and Di Pietro, 2008; Lopez-Berges et al., 2010). Briefly, cellophane membranes were cut the same size of a Petri dish, autoclaved in deionized water, and placed on top of PDA plates. 5 μl drops of in 2x10^7^ microconidia/ml in water were spot-inoculated at the center of the plate and plates were incubated at 28 °C for 3 d. After this time, the cellophane membrane with the fungal colony was carefully removed and the plates were incubated for another 24 h at 28 °C to visualize the mycelium that had penetrated through the cellophane. Plates were imaged before and after cellophane removal. All experiments were performed in triplicate.

### Plant infection assay

Tomato seeds (*Solanum lycopersicum* cv. Moneymaker from EELM-CSIC, or cv. Momotaro from Takii Seed Co., Ltd.; susceptible to *F. oxysporum* f. sp. *lycopersici* race 2) were surface-sterilized by immersion in 20% bleach (v/v) for 30 min and sown in moist vermiculite. Seedlings were grown in a growth chamber under the following conditions: 28 °C, 40-70% relative humidity and a photoperiod of 14 h of 36 W white light and 10 h of darkness.

Tomato plant infection assays were performed as described(Di Pietro and Roncero, 1998). Briefly, two-week-old seedlings were inoculated with the different fungal strains by immersing the roots in a suspension of 5x10^6^ microconidia/ml. Depending on the experiment, plants were irrigated with tap water or with a 10 µM CuSO_4_ solution. Disease symptoms and the survival rate were analyzed during 30-40 days. Death of the infected plants was diagnosed as a complete collapse of the stem, without any green parts left accompanied by visible proliferation of the fungal mycelium on the dead tissue. The Kaplan-Meier test was used to assess statistical significance of differences in survival among groups using the log-rank test with the software GraphPad Prims version v8.0.1(Lopez-Berges et al., 2012). All infection experiments were performed at least three times.

### Determination of *in planta* fungal burden

Fungal burden in tomato plants inoculated with *F. oxysporum* was measured by qPCR as described previously(Pareja-Jaime et al., 2010) using total DNA extracted from tomato roots and stems at 4 or 10 dpi. Relative fungal burden was calculated using the 2^-ΔΔCt^ method, with primers of the *Fol4287 six1* gene (*FOXG_16418*) and normalized to the tomato *gapdh* gene.

### Infection assays in *Galleria mellonella*

*G. mellonella* infection assays were performed as described(Navarro-Velasco et al., 2011; Navarro-Velasco et al., 2023). *G. mellonella* larvae (CASA REINA SA, Bilbao, Spain) were maintained in plastic boxes for 2-3 d before the infection. Fifteen larvae were used for each treatment. An automicroapplicator (0.1-10 μl; Burkard Manufacturing Co. Ltd) with a 1 ml syringe (Terumo Medical Corporation) was used to inject 8 μl of a 1.6x10^5^ microconidial suspension into the haemocoel of each larva. After injection, larvae were incubated in glass containers at 30 °C. Survival was recorded daily for 5 dpi. Data were analyzed with the software GraphPad Prims version v8.0.1(Lopez-Berges et al., 2012).

## Abbreviations

Mac1: <u>M</u>etal binding <u>ac</u>tivator <u>1</u>
Ctr: <u>C</u>opper <u>tr</u>ansport
Fre: <u>F</u>erric <u>re</u>ductase
Sod3: <u>S</u>uper<u>o</u>xide <u>d</u>ismutase <u>3</u>
Crm: <u>C</u>opper-responsive <u>m</u>etabolite

## Data availability

All raw quantitative data and statistical analyses are provided in the file Source Data. Uncropped PCR and Southern and Western blot images are provided in the file Raw Images. RNA-seq and ChIP-seq data have been deposited in the Gene Expression Omnibus database (Edgar et al., 2002) (Accession No. GSE243247 and GSE244102, respectively).

## Author Contributions and Acknowledgements

M.S.L.B., A.D.P., R.P.F., and J.S. designed the experiments. M.S.L.B., R.P.F., G.P.P., H.B. and L.S.R. carried out the experiments. M.S.L.B., A.D.P., R.P.F., M.V.A.P., G.P.P. and H.B. analyzed the data. M.S.L.B., A.D.P. and R.P.F. wrote the manuscript.

We are grateful to Mariló Alcaide Caballero and Florian Kastner for valuable technical assistance and to Rafael Fernández Muñoz, IHSM “La Mayora”, UMA-CSIC, Malaga, Spain, and Tsutomu Arie, Tokyo University of Agriculture and Technology, TUAT, Tokyo, Japan, for kindly providing seeds of tomato cultivars Moneymaker and Momotaro, respectively. This work was supported by grants from Junta de Andalucía (ProyExcel_00488) to M.S.L.B.; from the Spanish Ministry of Science and Innovation (MICINN, grant PID2022-140187OB-I00) to M.S.L.B. and A.D.P. and (MICINN, grants PLEC2021-007777, TED2021-130262B-I00 and PDC2022-133749-I00) to A.D.P.; and from the Austrian Science Fund (FWF, grant P32790-B) to J.S. and (FWF, grant T 1266) to L.S.R. R.P.F. was supported by Ph.D. fellowship FPU18/00028 from the Spanish Ministry of Universities and by the EMBO Scientific Exchange Grant 9756. M.V.A.P. was supported by the María Zambrano program to attract international talent 2021 from the Spanish Ministry of Universities and by an UCOLIDERA grant from University of Córdoba. G.P.P. was supported by Ph.D. fellowship FPI PRE2020-092679 from MICINN.

## References

Altschul, S.F., Gish, W., Miller, W., Myers, E.W., and Lipman, D.J. (1990). Basic local alignment search tool. Journal of Molecular Biology 215, 403–410.

Beaudoin, J., and Labbe, S. (2006). Copper Induces Cytoplasmic Retention of Fission Yeast Transcription Factor Cuf1. Eukaryot Cell 5, 277–292.

Berendsen, R.L., Pieterse, C.M., and Bakker, P.A. (2012). The rhizosphere microbiome and plant health. Trends Plant Sci 17, 478–486.

Bolger, A.M., Lohse, M., and Usadel, B. (2014). Trimmomatic: a flexible trimmer for Illumina sequence data. Bioinformatics 30, 2114–2120.

Cai, Z., Du, W., Liu, L., Pan, D., and Lu, L. (2019). Molecular Characteristics of the Conserved *Aspergillus nidulans* Transcription Factor Mac1 and Its Functions in Response to Copper Starvation. mSphere 4.

Cai, Z., Du, W., Zeng, Q., Long, N., Dai, C., and Lu, L. (2017). Cu-sensing transcription factor Mac1 coordinates with the Ctr transporter family to regulate Cu acquisition and virulence in *Aspergillus fumigatus*. Fungal genetics and biology : FG & B 107, 31–43.

Culbertson, E.M., Bruno, V.M., Cormack, B.P., and Culotta, V.C. (2020). Expanded role of the Cu-sensing transcription factor Mac1p in *Candida albicans*. Mol Microbiol 114, 1006–1018.

Dean, R., Van Kan, J.A.L., Pretorius, Z.A., Hammond-Kosack, K.E., Di Pietro, A., Spanu, P.D., Rudd, J.J., Dickman, M., Kahmann, R., Ellis, J., and Foster, G.D. (2012). The Top 10 fungal pathogens in molecular plant pathology. Mol. Plant Pathol. 13, 414–430.

Di Pietro, A., and Roncero, M.I. (1998). Cloning, expression, and role in pathogenicity of *pg1* encoding the major extracellular endopolygalacturonase of the vascular wilt pathogen *Fusarium oxysporum*. Mol Plant Microbe Interact. 11, 91–98.

Edgar, R., Domrachev, M., and Lash, A.E. (2002). Gene Expression Omnibus: NCBI gene expression and hybridization array data repository. Nucleic Acids Res 30, 207–210.

Fernandes, T.R., Segorbe, D., Prusky, D., and Di Pietro, A. (2017). How alkalinization drives fungal pathogenicity. PLoS Pathog 13, e1006621.

Festa, R.A., and Thiele, D.J. (2011). Copper: an essential metal in biology. Current biology : CB 21, R877–883.

Gacek-Matthews, A., Berger, H., Sasaki, T., Wittstein, K., Gruber, C., Lewis, Z.A., and Strauss, J. (2016). KdmB, a Jumonji Histone H3 Demethylase, Regulates Genome-Wide H3K4 Trimethylation and Is Required for Normal Induction of Secondary Metabolism in *Aspergillus nidulans*. PLoS Genet 12, e1006222.

Gel, B., and Serra, E. (2017). karyoploteR: an R/Bioconductor package to plot customizable genomes displaying arbitrary data. Bioinformatics 33, 3088–3090.

Goswami, R.S. (2012). Targeted gene replacement in fungi using a split-marker approach. Methods in molecular biology 835, 255–269.

Grant, C.E., Bailey, T.L., and Noble, W.S. (2011). FIMO: scanning for occurrences of a given motif. Bioinformatics 27, 1017–1018.

Gu, Z., Eils, R., and Schlesner, M. (2016). Complex heatmaps reveal patterns and correlations in multidimensional genomic data. Bioinformatics 32, 2847–2849.

Hervas-Aguilar, A., and Penalva, M.A. (2010). Endocytic machinery protein SlaB is dispensable for polarity establishment but necessary for polarity maintenance in hyphal tip cells of *Aspergillus nidulans*. Eukaryot Cell 9, 1504–1518.

Jungmann, J., Reins, H.A., Lee, J., Romeo, A., Hassett, R., Kosman, D., and Jentsch, S. (1993). MAC1, a nuclear regulatory protein related to Cu-dependent transcription factors is involved in Cu/Fe utilization and stress resistance in yeast. The EMBO journal 12, 5051**-**5056.

Katoh, K., and Standley, D.M. (2013). MAFFT multiple sequence alignment software version 7: improvements in performance and usability. Mol Biol Evol 30, 772–780.

Kopke, K., Hoff, B., and Kuck, U. (2010). Application of the *Saccharomyces cerevisiae* FLP/FRT recombination system in filamentous fungi for marker recycling and construction of knockout strains devoid of heterologous genes. Appl Environ Microbiol 76, 4664–4674.

Krassowski, M. (2020). ComplexUpset.

Lex, A., Gehlenborg, N., Strobelt, H., Vuillemot, R., and Pfister, H. (2014). UpSet: Visualization of Intersecting Sets. IEEE transactions on visualization and computer graphics 20, 1983–1992.

Li, H., and Durbin, R. (2009). Fast and accurate short read alignment with Burrows– Wheeler transform. Bioinformatics 25, 1754–1760.

Lim, F.Y., Won, T.H., Raffa, N., Baccile, J.A., Wisecaver, J., Rokas, A., Schroeder, F.C., and Keller, N.P. (2018). Fungal Isocyanide Synthases and Xanthocillin Biosynthesis in *Aspergillus fumigatus*. mBio 9.

Livak, K.J., and Schmittgen, T.D. (2001). Analysis of relative gene expression data using real-time quantitative PCR and the 2(-Delta Delta C(T)) Method. Methods (San Diego, Calif.) 25, 402–408.

Lopez-Berges, M.S., Rispail, N., Prados-Rosales, R.C., and Di Pietro, A. (2010). A nitrogen response pathway regulates virulence functions in *Fusarium oxysporum* via the protein kinase TOR and the bZIP protein MeaB. Plant Cell 22, 2459–2475.

Lopez-Berges, M.S., Pinar, M., Abenza, J.F., Arst, H.N., Jr., and Penalva, M.A. (2016). The *Aspergillus nidulans* syntaxin PepA(Pep12) is regulated by two Sec1/Munc-18 proteins to mediate fusion events at early endosomes, late endosomes and vacuoles. Mol Microbiol 99, 199–216.

Lopez-Berges, M.S., Capilla, J., Turra, D., Schafferer, L., Matthijs, S., Jochl, C., Cornelis, P., Guarro, J., Haas, H., and Di Pietro, A. (2012). HapX-mediated iron homeostasis is essential for rhizosphere competence and virulence of the soilborne pathogen *Fusarium oxysporum*. Plant Cell 24, 3805–3822.

Lopez-Berges, M.S., Scheven, M.T., Hortschansky, P., Misslinger, M., Baldin, C., Gsaller, F., Werner, E.R., Kruger, T., Kniemeyer, O., Weber, J., Brakhage, A.A., and Haas, H. (2021). The bZIP Transcription Factor HapX Is Post-Translationally Regulated to Control Iron Homeostasis in *Aspergillus fumigatus*. International journal of molecular sciences 22.

López-Berges, M.S. (2020). ZafA-mediated regulation of zinc homeostasis is required for virulence in the plant pathogen *Fusarium oxysporum*. Molecular Plant Pathology 21, 244–249.

López-Berges, M.S., Capilla, J., Turrà, D., Schafferer, L., Matthijs, S., Joechl, C., Cornelis, P., Guarro, J., Haas, H., and Di Pietro, A. (2012). HapX-mediated iron homeostasis is essential for rhizosphere competence and virulence of the soilborne pathogen *Fusarium oxysporum*. Plant Cell 24, 3805–3822.

Love, M.I., Huber, W., and Anders, S. (2014). Moderated estimation of fold change and dispersion for RNA-seq data with DESeq2. Genome biology 15, 550.

Mattern, I.E., Punt, P.J., and Van den Hondel, C.A. (1988). A vector for Aspergillus transformation conferring phleomycin resistance. Fungal Genetics Reports 35.

Navarro-Velasco, G.Y., Di Pietro, A., and Lopez-Berges, M.S. (2023). Constitutive activation of TORC1 signalling attenuates virulence in the cross-kingdom fungal pathogen *Fusarium oxysporum*. Mol Plant Pathol 24, 289–301.

Navarro-Velasco, G.Y., Prados-Rosales, R.C., Ortiz-Urquiza, A., Quesada-Moraga, E., and Di Pietro, A. (2011). *Galleria mellonella* as model host for the trans-kingdom pathogen *Fusarium oxysporum*. Fungal genetics and biology : FG & B 48, 1124–1129.

Nucci, M., and Anaissie, E. (2007). *Fusarium* infections in immunocompromised patients. Clinical Microbiology Reviews 20, 695–704.

Ordonez, N., Seidl, M.F., Waalwijk, C., Drenth, A., Kilian, A., Thomma, B.P., Ploetz, R.C., and Kema, G.H. (2015). Worse Comes to Worst: Bananas and Panama Disease--When Plant and Pathogen Clones Meet. PLoS Pathog 11, e1005197.

Pareja-Jaime, Y., Martin-Urdiroz, M., Roncero, M.I., Gonzalez-Reyes, J.A., and Ruiz-Roldan, M.C. (2010). Chitin synthase deficient mutant of Fusarium oxysporum elicits tomato plant defence response and protects against wild type infection. Mol Plant Pathol. *in press*.

Park, Y.-S., Lian, H., Chang, M., Kang, C.-M., and Yun, C.-W. (2014). Identification of high-affinity copper transporters in *Aspergillus fumigatus*. Fungal Genetics and Biology 73, 29–38.

Park, Y.S., Kim, T.H., and Yun, C.W. (2017). Functional characterization of the copper transcription factor AfMac1 from *Aspergillus fumigatus*. The Biochemical journal 474, 2365–2378.

Park, Y.S., Kang, S., Seo, H., and Yun, C.W. (2018). A copper transcription factor, AfMac1, regulates both iron and copper homeostasis in the opportunistic fungal pathogen *Aspergillus fumigatus*. The Biochemical journal 475, 2831–2845.

Patro, R., Duggal, G., Love, M.I., Irizarry, R.A., and Kingsford, C. (2017). Salmon provides fast and bias-aware quantification of transcript expression. Nature methods 14, 417–419.

Pena, M.M., Puig, S., and Thiele, D.J. (2000). Characterization of the *Saccharomyces cerevisiae* high affinity copper transporter Ctr3. The Journal of biological chemistry 275, 33244–33251.

Pfaffl, M.W. (2001). A new mathematical model for relative quantification in real-time RT-PCR. Nucleic acids research 29, e45.

Pinar, M., Arst, H.N., Jr., Pantazopoulou, A., Tagua, V.G., de los Rios, V., Rodriguez-Salarichs, J., Diaz, J.F., and Penalva, M.A. (2015). TRAPPII regulates exocytic Golgi exit by mediating nucleotide exchange on the Ypt31 ortholog RabERAB11. Proc Natl Acad Sci U S A 112, 4346–4351.

Prados Rosales, R.C., and Di Pietro, A. (2008). Vegetative hyphal fusion is not essential for plant infection by *Fusarium oxysporum*. Eukaryot Cell 7, 162–171.

Punt, P.J., Oliver, R.P., Dingemanse, M.A., Pouwels, P.H., and van den Hondel, C.A. (1987). Transformation of Aspergillus based on the hygromycin B resistance marker from Escherichia coli. Gene 56, 117–124.

Qi, J., Han, A., Yang, Z., and Li, C. (2012). Metal-sensing transcription factors Mac1p and Aft1p coordinately regulate vacuolar copper transporter CTR2 in *Saccharomyces cerevisiae*. Biochem Biophys Res Commun 423, 424–428.

Raffa, N., Osherov, N., and Keller, N.P. (2019). Copper Utilization, Regulation, and Acquisition by *Aspergillus fumigatus*. International journal of molecular sciences 20.

Ray, S.C., and Rappleye, C.A. (2022). Mac1-Dependent Copper Sensing Promotes *Histoplasma* Adaptation to the Phagosome during Adaptive Immunity. mBio 13, e0377321.

Redkar, A., Sabale, M., Schudoma, C., Zechmann, B., Gupta, Y.K., Lopez-Berges, M.S., Venturini, G., Gimenez-Ibanez, S., Turra, D., Solano, R., and Di Pietro, A. (2022). Conserved secreted effectors contribute to endophytic growth and multihost plant compatibility in a vascular wilt fungus. Plant Cell 34, 3214–3232.

Reyes-Dominguez, Y., Boedi, S., Sulyok, M., Wiesenberger, G., Stoppacher, N., Krska, R., and Strauss, J. (2012). Heterochromatin influences the secondary metabolite profile in the plant pathogen *Fusarium graminearum*. Fungal genetics and biology : FG & B 49, 39–47.

Rutherford, J.C., and Bird, A.J. (2004). Metal-responsive transcription factors that regulate iron, zinc, and copper homeostasis in eukaryotic cells. Eukaryot Cell 3, 1–13.

Ryan, B.M., Kirby, J.K., Degryse, F., Harris, H., McLaughlin, M.J., and Scheiderich, K. (2013). Copper speciation and isotopic fractionation in plants: uptake and translocation mechanisms. The New phytologist 199, 367–378.

Schneider-Poetsch, T., Ju, J., Eyler, D.E., Dang, Y., Bhat, S., Merrick, W.C., Green, R., Shen, B., and Liu, J.O. (2010). Inhibition of eukaryotic translation elongation by cycloheximide and lactimidomycin. Nat Chem Biol 6, 209–217.

Schrettl, M., Beckmann, N., Varga, J., Heinekamp, T., Jacobsen, I.D., Joechl, C., Moussa, T.A., Wang, S., Gsaller, F., Blatzer, M., Werner, E.R., Niermann, W.C., Brakhage, A.A., and Haas, H. (2010). HapX-Mediated Adaption to Iron Starvation Is Crucial for Virulence of *Aspergillus fumigatus*. Plos Pathogens 6.

Shi, H., Jiang, Y., Yang, Y., Peng, Y., and Li, C. (2021). Copper metabolism in *Saccharomyces cerevisiae*: an update. Biometals 34, 3–14.

Smith, A.D., Logeman, B.L., and Thiele, D.J. (2017). Copper Acquisition and Utilization in Fungi. Annu Rev Microbiol 71, 597–623.

Sun, T.S., Ju, X., Gao, H.L., Wang, T., Thiele, D.J., Li, J.Y., Wang, Z.Y., and Ding, C. (2014). Reciprocal functions of *Cryptococcus neoformans* copper homeostasis machinery during pulmonary infection and meningoencephalitis. Nature communications 5, 5550.

Tamura, K., Stecher, G., and Kumar, S. (2021). MEGA11: Molecular Evolutionary Genetics Analysis Version 11. Mol Biol Evol 38, 3022–3027.

Team, R.C. (2023). R: A language and environment for statistical computing (Versión, 4.3.0). R Foundation for Statistical Computing.

Torres, A.M., Weeden, N.F., and Martín, A. (1993). Linkage among isozyme, RFLP and RAPD markers in Vicia faba. Theoretical and applied genetics 85, 937–945.

Turra, D., El Ghalid, M., Rossi, F., and Di Pietro, A. (2015). Fungal pathogen uses sex pheromone receptor for chemotropic sensing of host plant signals. Nature 527, 521–524.

Waterman, S.R., Hacham, M., Hu, G., Zhu, X., Park, Y.D., Shin, S., Panepinto, J., Valyi-Nagy, T., Beam, C., Husain, S., Singh, N., and Williamson, P.R. (2007). Role of a CUF1/CTR4 copper regulatory axis in the virulence of *Cryptococcus neoformans*. The Journal of clinical investigation 117, 794–802.

Wegner, S.V., Sun, F., Hernandez, N., and He, C. (2011). The tightly regulated copper window in yeast. Chemical communications 47, 2571–2573.

Wei, T., and Simko, V. (2021). R package ‘corrplot’: Visualization of a Correlation Matrix.

Wiemann, P., Perevitsky, A., Lim, F.Y., Shadkchan, Y., Knox, B.P., Landero Figueora, J.A., Choera, T., Niu, M., Steinberger, A.J., Wuthrich, M., Idol, R.A., Klein, B.S., Dinauer, M.C., Huttenlocher, A., Osherov, N., and Keller, N.P. (2017). *Aspergillus fumigatus* Copper Export Machinery and Reactive Oxygen Intermediate Defense Counter Host Copper-Mediated Oxidative Antimicrobial Offense. Cell reports 19, 2174–2176.

Won, T.H., Bok, J.W., Nadig, N., Venkatesh, N., Nickles, G., Greco, C., Lim, F.Y., Gonzalez, J.B., Turgeon, B.G., Keller, N.P., and Schroeder, F.C. (2022). Copper starvation induces antimicrobial isocyanide integrated into two distinct biosynthetic pathways in fungi. Nature communications 13, 4828.

Zhu, Z., Labbe, S., Pena, M.M., and Thiele, D.J. (1998). Copper differentially regulates the activity and degradation of yeast Mac1 transcription factor. The Journal of biological chemistry 273, 1277–1280.

